# Evidence based Unification of poTato gene models with UniTato collaborative genome browser

**DOI:** 10.1101/2023.12.07.570586

**Authors:** Maja Zagorščak, Jan Zrimec, Carissa Bleker, Nadja Nolte, Živa Ramšak, Kristina Gruden, Marko Petek

**Affiliations:** National Institute of Biology, Večna pot 121, 1000 Ljubljana, Slovenia

## Abstract

Potato (*Solanum tuberosum*) is the most popular tuber crop and model organism. Though its gene models are frequently updated, they are not unified, leading to missing or wrongly annotated genes. Here, we thus unify the recent potato double monoploid v4 and v6 gene models by automatic merging. We established an Apollo web server that enables access to the Unified poTato genome annotation database (UniTato) as well as further community-based manual curation at <u>unitato.nib.si</u>. We demonstrate how the web server interface can help to resolve problems with missing or misplaced genes and can be used to further update them or consolidate a wider set of gene models or genome information. Genome annotation files and a comprehensive translation table are provided at github.com/NIB-SI/unitato.

## 1. Introduction

*Solanum tuberosum* (potato) is among the most important food crops and a model tuber species. Apart from novel wild potato (Tang et al., 2022) and pan-genome assemblies (Hoopes et al., 2022; Bozan et al., 2023), the double monoploid (DM) clone of group Phureja DM1-3 516 R44 was until recently the standard variety on which gene models were defined. Multiple DM genome assemblies and gene models versions have been introduced by different consortia, including the Potato Genome Sequencing Consortium (PGSC), International Tomato Genome Consortium (ITAG) and Buell Lab (University of Georgia). These sequenced and assembled (Yandell and Ence, 2012) up to ∼88% of the potato genome that included from 35,004 (ITAG) (Tomato Genome Consortium, 2012) to 39,428 (PGSCv4.04) (Potato Genome Sequencing Consortium et al., 2011) gene models, with the recent nanopore long read assembly DMv6.1 (Pham et al., 2020) annotating 40,652 genes (in the working version). Moreover, we previously unified the v4 PGSC and ITAG gene models into a merged DMv4n version containing 49,322 genes (Petek et al., 2020).

Despite progressive assembly improvements, we observed that the recent v6 gene models did not include certain known genes with important molecular functions, as they do not account for previous gene model information. An example is the transcription factor TGA2, an essential regulator of hormonal signalling (Tomaž et al., 2023). Aside from such missing genes, some are also moved, merged or split, which can lead to differences in interpretation in downstream analyses (e.g. gene family expansion, differential expression, marker selection, gene set enrichment analysis) (Yandell and Ence, 2012). In addition to these incomplete annotations negatively affecting further experiments, existing published results using previous gene model versions, including e.g. AlphaFold structure predictions (Tomaž et al., 2023), have become outdated, making it essential to update and consolidate gene predictions.

To help resolve these issues, we expand the ITAG and PGSC v4 annotations with v6 annotations (Pham et al., 2020), integrating and unifying the different gene models. In addition, we include experiment-based evidence from our pan-transcriptome (Petek et al., 2020), thereby creating an improved and more accurate potato gene model annotation for downstream analyses. In order to ensure the transparency and accuracy of future gene models, we present the Unified poTato genome annotation resource (UniTato), hosted on an Apollo web server (Dunn et al., 2019), which enables a community-driven effort for real-time revision and enhancement of gene models by experts. This will increase interpretational power of experimental datasets and facilitate the reuse of experimental analyses conducted on v4 and expedite further progress in potato research.

## 2. Methods

### 2.1 Data sources

To develop the database, we used publicly available potato Group Phureja DM gene models: DMv6.1 (Pham et al., 2020), ITAG (Tomato Genome Consortium, 2012), and PGSCv4.04 (Potato Genome Sequencing Consortium et al., 2011), as well as reference transcriptomes of Désirée, Rywal, and PW363 tetraploid genotypes, and ITAG/PGSC translation table (Petek et al., 2020). The latter consolidated the two publicly available PGSC and ITAG gene models into a single unified one.

### 2.2 Data processing

To map gene annotations across potato genome assemblies (Figure 1), GFF files were sorted using the *sort* function from Bedtools v2.25.0 (Quinlan and Hall, 2010). Liftoff v.1.6.3 (Shumate and Salzberg, 2020) uses Minimap2 (Li, 2018) to map annotations between assemblies of the same or closely related species. We modified it to accept the number of nucleotides for the *flank* parameter (https://github.com/NIB-SI/Liftoff), instead of the ratio of sequence size, and used with the following parameters: (i) coverage of 0.90%, (ii) sequence identity of 90%, (iii) flanking sequence length *flank* of either 0 or 500 nucleotides, and (iv) Minimap2 v.2.24-r1122 ‘asm5’ option for long assembly to reference mapping. In addition, Minimap2 was used with the same set of parameters as for Liftoff (--end-bonus 5 --eqx -N 50 -p 0.8 -ax asm5) to map the reference CDSome and transcriptome (Petek et al., 2020) of three potato genotypes: Désirée, PW363 and Rywal. FASTQ files of long read transcriptomics datasets were downloaded from SRA. The Iso-Seq reads were mapped to the v6 genome assembly using minimap2 with parameters “-ax splice:hq -G 10000 -uf”. Next, to compare the mapped annotation across the assemblies, as well as overlaps within the DMv6.1, the *intersect* function from Bedtools (Quinlan and Hall, 2010) was used with the following minimum overlap as a fraction (*F*) ranging incrementally from 0.0001 to 1. Pairs from our existing merged v4 genome model (Petek et al., 2020) were used to determine the optimal *F* threshold value of 0.30 (Supp. figure S1). All reported v6 values below refer to working model versions unless stated otherwise. For visualisation, packages circlize v0.4.15 (Gu et al., 2014) and intervals v0.15.4 (github.com/edzer/intervals) were used with default settings. For topological sorting of unified gff features, AGAT v0.6.2 (Dainat et al., 2023) was used with default settings. The programming environments R v.4.3 (https://www.r-project.org/) and Python v3.8 (https://www.python.org/) were used. Code to reproduce the analysis and results including scripts used for constructing the mapping table between v4 and v6 gene IDs, as well as merging v4 and v6 models are available at the GitHub repository https://github.com/NIB-SI/unitato.

**Figure 1.**
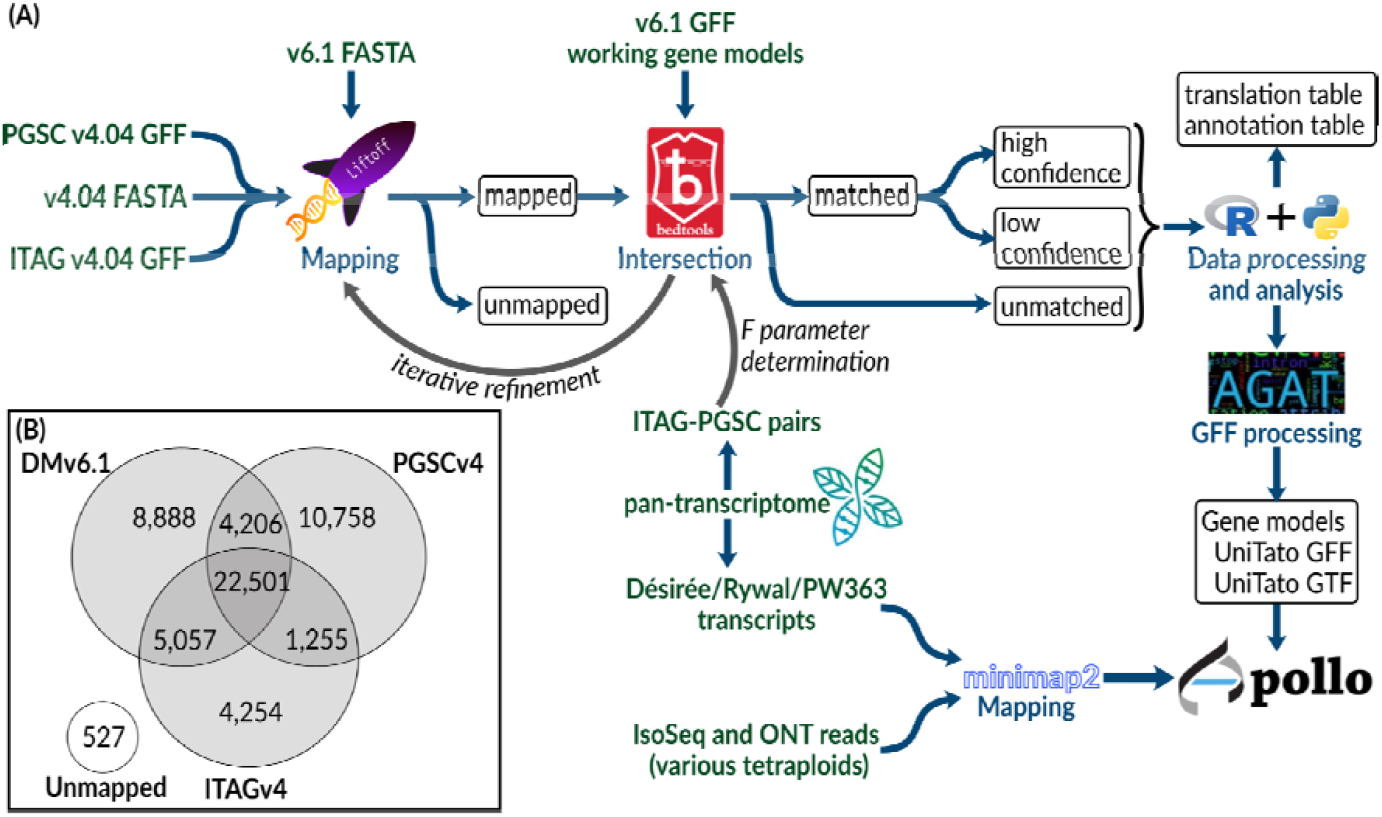
Setting up UniTato potato DM gene models. (A) Schematic overview of the procedure used to create a unified DM v4 and v6 potato genome annotation resource. (B) Venn diagram of resulting overlaps of the different v4 and v6 gene models (F > 0.30). Note that for intersected areas v6 IDs are shown.

### 2.3 Database implementation

A web server hosting the Apollo genomic annotation editor (Dunn et al., 2019) for real-time collaborative analysis and curation was deployed at https://unitato.nib.si. The reference DM genome assembly (DMv6.1) was uploaded as the base organism. Several evidence tracks corresponding to different gene models are available for exploration and curation (detailed in Figure 3A). The database instance is running Apollo 2.7.0, deployed with docker, with default parameters. Data upload was carried out using JBrowse utility scripts (Buels et al., 2016).

## 3. Results

### 3.1 A unified v4 and v6 potato genome annotation

To compare the potato gene model versions, we mapped gene annotations of older PGSCv4 (Potato Genome Sequencing Consortium et al., 2011) and ITAG assemblies (Tomato Genome Consortium, 2012) to the recent potato DMv6.1 assembly (Pham et al., 2020) using Liftoff (Shumate and Salzberg, 2020) and used Bedtools *intersect* (Quinlan and Hall, 2010) to find intersecting genes (Figure 1, see Methods 2.2). Briefly, Liftoff is a tool that accurately maps annotations between assemblies of the same or closely related species. We used it to transfer the gene model annotations from v4 to the v6 assembly. Two genome assemblies (either v4 PGSC or ITAG and DMv6.1) and a v4 annotation file (v4 PGSC or ITAG, respectively) were provided as input. The v4 gene models were aligned chromosome by chromosome to the v6 genome assembly. Here, a key parameter termed *flank* controls the amount of flanking sequence upstream and downstream of a gene. Bedtools *intersect* (Quinlan and Hall, 2010) was then used to check for overlap (intersection) between the sets of v4 and v6 gene models. The parameter *F* allows for control over the minimum overlap required as a fraction of the v4 gene models.

We first explored the Liftoff *flank* parameter, using either none or 500 nt of flanking sequence. The upstream and downstream expansion of each v4 gene sequence (combined PGSC/ITAG v4 dataset), before mapping, to also include the neighbourhood of the gene, generally improves mapping precision. This is especially important for the ITAG annotation which contains only CDS regions, whereas with the PGSC annotation, complete mRNA sequences are provided. Without a flanking sequence, we mapped 72,143 v4 gene models, whereas using a flanking sequence of length 500 nt we mapped 73,820 v4 gene models. Using both flank parameter values, 316 PGSC and 211 ITAG gene models could not be mapped to the v6 genome assembly (Supp. table S1).

We next explored the Bedtools *F* parameter, relaying the coverage of v4 sequence mapping to v6 and ranging from 0.0001 to 1, and found that 0.30 was the optimum value (Supp. figure S1, Supp. table S2). Setting the Liftoff flanking parameter to 500 nt, we achieved a mapping coverage (*F* >= 0.3, high identity) with 56,776 v4 gene models mapping to 31,594 v6 models (of these, 92% belong to v6 high confidence gene models, Supp. table S3). Since *flank* can also capture v4 assembly gaps (N runs) or misassemblies that were corrected in the v6 assembly, using it may not always be the optimal choice. For example, we found that the lack of a flanking sequence (0 nt) achieved a better mapping coverage *F* with 387 v4 gene models mapping to 458 v6 models. We thus decided to keep the Liftoff result with the better mapping coverage per gene (either 0 or 500 nt *flank)*, as reported above. For gene models with a Bedtools coverage *F* above or equal to 0.30, we kept the v6 gene models and added 17,272 v4 models with low coverage (*F* < 0.30). This merge resulted in the genome annotation termed UniTato (Figure 2). Note that the v6 genome assembly has many inversions compared to v4 assembly, most evidently in chromosome 12 (Supp. figure S3).

**Figure 2.**
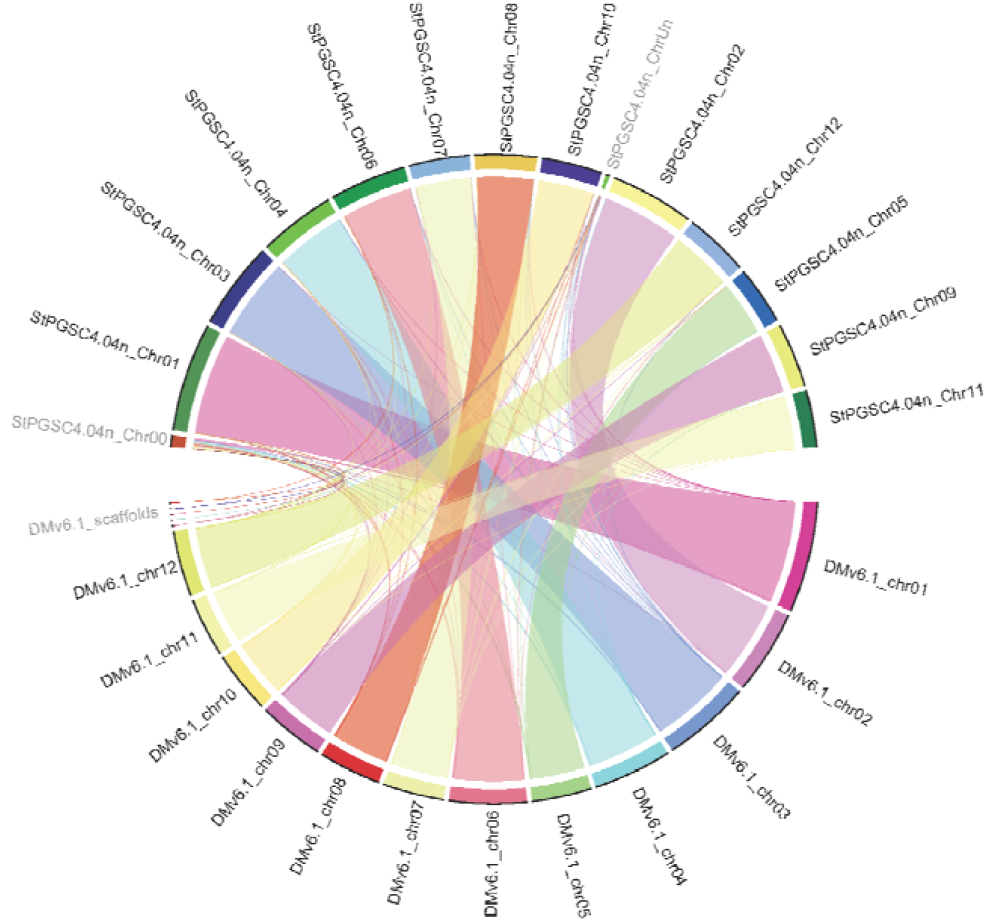
Chord diagram of the synteny between v4 and v6 gene models. The diagram shows that most chromosomes are almost completely syntenic across models, however some scaffolds remain unanchored.

**Figure 3.**
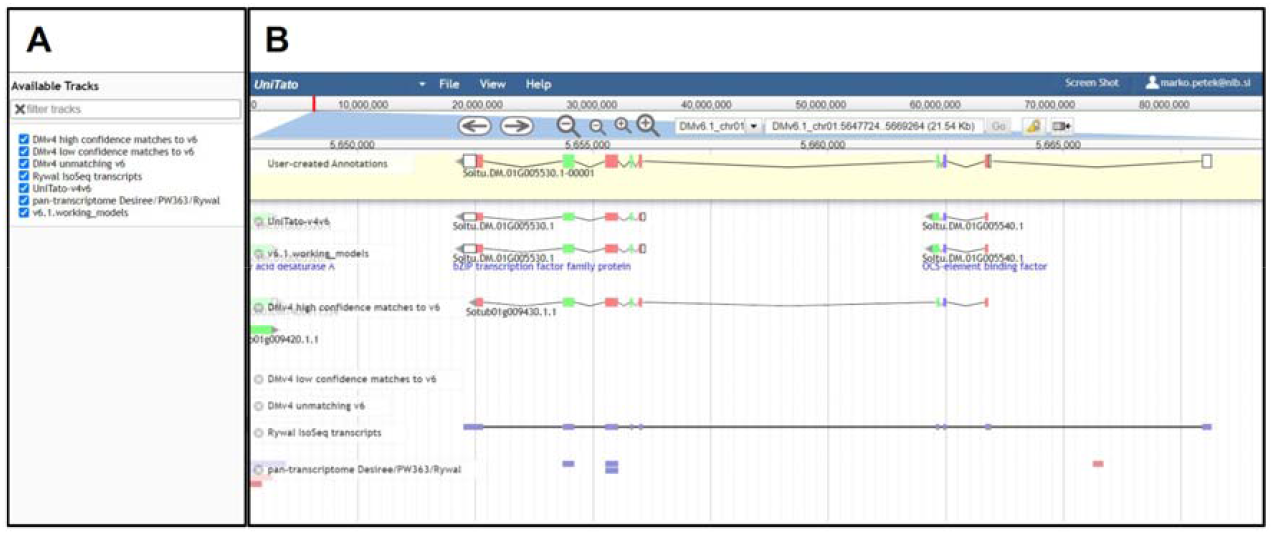
Overview of the UniTato user interface and TGA2 use case. Screenshot of the Apollo server web interface for the *Solanum tuberosum* DM gene model manual annotation. (A) Tracks with annotations to aid manual corrections. (B) Manual annotation of a TGA2 transcription factor gene model which was split into two gene models in v6 (track v6.1.working_models). The gene model’s manual annotation with nine exons (track “User-created Annotations”) was based on the correctly predicted ITAG v4 CDS and the Rywal Iso-Seq transcript mapping.

Of the observed 17,272 v4 gene models with low coverage (*F* < 0.30), 11,832 were from the PGSC dataset and 5,440 from ITAG. These sequences are present in the v6 assembly but were not identified as genes (Pham et al., 2020). We decided to retain all such “rescued” genes and assigned them with the identifier from v4. Of these, 16,117 mapped to the intergenic regions in v6 (*F* < 0.0001). On the other hand, 7,381 v6 working version gene models were not supported by v4 annotations (of these, 5,590 with v6 annotation ‘hypothetical protein’), of which 2,673 were high-confidence v6 gene models. Finally, we further analysed the genome-mapped and unmapped v4 genes, searching for evidence of their expression within our published pan-transcriptome dataset (Petek et al., 2020). The v4 gene models that do not match any v6 gene models (*F* < 0.0001), but do match tetraploid transcriptomes (3,596 out of 15,590 gene models) are considered valid genes. On the other hand, the 11,924 that match neither the v6 models nor the pan-transcriptome are likely unreliable gene model predictions. Note that 292 out of 559 v4 gene models did not map to the v6 genome yet match tetraploid Désirée, Rywal, or PW363 transcripts. These genes were lost with reassembly of the DM scaffold in v6 (Pham et al., 2020). The newly generated GFF3 file and a table linking identifiers of PGSC, iTAG v4 gene models with v6 version is available at GitHub (https://github.com/NIB-SI/unitato).

### 3.2 UniTato database access and user interface

The UniTato database (accessible at http://unitato.nib.si/) is hosted in a deployment of the community-focused genome annotation editor Apollo (Dunn et al., 2019). Based on the popular JBrowse genome viewer (Buels et al., 2016), Apollo allows visitors to browse, compare, and interpret the available evidence-based gene models. The annotator panel in the Apollo interface provides several tabs, allowing easy navigation through the genome, the ability to view or hide tracks, as well as to locate and view annotation details. For further information, refer to the Apollo online documentation (https://genomearchitect.readthedocs.io/).

The database currently contains a number of evidence tracks, including (i) v6 working gene models, (ii) Rywal mapped Iso-Seq reads, (iii) unified v4 and v6 potato genome annotation (UniTato) (iv) v4 gene models that mapped to the v6 intergenic regions (DMv4 unmatching v6), (v) low and (vi) high confidence v4 gene model matches to v6 models, and (vii) mapped tetraploid potato transcripts (pan-transcriptome Désirée/PW363/Rywal). These evidence tracks are publicly viewable by all UniTato web page visitors (Figure 3A). On the other hand, potential contributors are encouraged to use the contact details on the web page to request edit access through a user account. Upon login, these users have access to the curator tools, providing the ability to collaboratively add, remove and modify potato gene models. The improvements can then be exported as a new version of the genome annotation file (GFF, VCF, or FASTA).

### 3.3 UniTato improves the coverage and accuracy of gene models

Merging of v4 and v6 genome annotations improves the coverage and accuracy of gene models, whereas manual annotation by the experts will provide the necessary quality control to further improve the accuracy of gene models. The improved coverage is most evident by adding the rescued v4 genes showing experimental evidence for expression. These include certain important genes, such as a gene encoding a cysteine protease inhibitor (PGSC0003DMG400010139/Sotub03g015980) and the salicylic acid-binding protein 2 (PGSC0003DMG400028777/Sotub06g025780; for details see v4-v6.1_translationTable.xlsx on GitHub). Apart from the missing genes, several v6 genome models have been wrongly predicted. One such case is the TGA2 transcription factor gene encoded by two v6 gene models and correctly annotated as a single gene model by ITAG v4 (Tomaž et al., 2023). The Iso-Seq read mapping suggests that the gene’s 5’-untranslated region extends into another exon (Figure 3B). Such misannotations can be easily manually curated in UniTato where tracks of mapped transcripts can help manual curators to build more accurate gene models. Furthermore, we observed several more complicated cases of gene models that cannot be simply resolved between v4 and v6 annotations and will need to be manually curated (see example in Supp. figure S2). We also provide a list of gene identifiers for these complicated cases on GitHub (see overlaps.xlsx on GitHub).

## 4. Discussion

The advancement and maturation of high-throughput and long-read sequencing has led to a number of different potato genome assemblies, gene annotations and transcriptomics datasets. Sequencing the group Phureja DM (Potato Genome Sequencing Consortium et al., 2011) still enables functional studies of polyploid potato cultivars using RNA-Seq technologies, although with the limitation of not covering cultivar-specific gene expression (Petek et al., 2020; Hoopes et al., 2022). Most potato researchers use only one genome annotation, either PGSC (Potato Genome Sequencing Consortium et al., 2011) or ITAG (Tomato Genome Consortium, 2012), for practical reasons, especially when conducting high-throughput analyses. However, using an incomplete gene set can lead to false outcomes regarding gene presence or gene family diversity, severely affecting downstream results (Yandell and Ence, 2012; Petek et al., 2020). It is well known that incorrect or incomplete annotations corrupt all subsequent experiments that rely on them, making it absolutely essential to have the ability to provide accurate and up-to-date annotations (Yandell and Ence, 2012; Bolger et al., 2018).

Our motivation here was thus two-fold: first, to transfer both gene model sets from the older PGSC assembly (Potato Genome Sequencing Consortium et al., 2011; Tomato Genome Consortium, 2012) to the newly upgraded one (Pham et al., 2020) and, at the same time, to merge the gene models, allowing for data interoperability of previous experimental results (e.g. from RNA-Seq) (Petek et al., 2020) with the unified gene model set, UniTato. Annotation merging was performed using an in-house developed bioinformatics pipeline; utilising open-source software and complementing it with the evidence from published tetraploid transcriptomes (Petek et al., 2020). The resulting annotation files were incorporated in an Apollo web server (Dunn et al., 2019), which enables the potato community to curate and refine potato gene models on-site and in real-time, facilitating the establishment of a single standardised potato genome annotation.

The requirements of periodic annotation curation and incorporating experimental data and novel findings into the annotation process are inherent also to other plant species (Yandell and Ence, 2012; Kersey, 2019). We propose that a similar approach for evidence and community-based revision as the one presented here can be applied also with those organisms. Apart from updating our database with new assemblies as they become available (Yang et al., 2023), future developments include the addition of novel experimental omics datasets and possibly expansion to related genomes.

## Data availability statement

Data and code to reproduce the analysis are available at the GitHub repository https://github.com/NIB-SI/unitato/.

## Author contributions

MZ, MP and KG conceptualised the project. MZ, JZ, KG and MP designed the computational pipelines. MZ, CB and NN performed the computational analysis. MZ, JZ, MP and KG interpreted the results. MZ, JZ, CB and MP wrote the draft and all authors contributed to the final manuscript.

## References

Bolger, M. E., Arsova, B., and Usadel, B. (2018). Plant genome and transcriptome annotations: from misconceptions to simple solutions. Brief. Bioinform. 19, 437–449.

Bozan, I., Achakkagari, S. R., Anglin, N. L., Ellis, D., Tai, H. H., and Strömvik, M. V. (2023). Pangenome analyses reveal impact of transposable elements and ploidy on the evolution of potato species. Proc. Natl. Acad. Sci. U. S. A. 120, e2211117120.

Buels, R., Yao, E., Diesh, C. M., Hayes, R. D., Munoz-Torres, M., Helt, G., et al. (2016). JBrowse: a dynamic web platform for genome visualization and analysis. Genome Biol. 17, 66.

Dainat, J., Hereñú D., Murray, K. D., Davis E., Crouch, K., Sol, L., Agostinho, N., et al. (2023). NBISweden/AGAT: AGAT-v1.2.0. Zenodo. doi: 10.5281/zenodo.8178877.

Dunn, N. A., Unni, D. R., Diesh, C., Munoz-Torres, M., Harris, N. L., Yao, E., et al. (2019). Apollo: Democratizing genome annotation. PLoS Comput. Biol. 15, e1006790.

Gu, Z., Gu, L., Eils, R., Schlesner, M., and Brors, B. (2014). circlize Implements and enhances circular visualization in R. Bioinformatics 30, 2811–2812.

Hoopes, G., Meng, X., Hamilton, J. P., Achakkagari, S. R., de Alves Freitas Guesdes, F., Bolger, M. E., et al. (2022). Phased, chromosome-scale genome assemblies of tetraploid potato reveal a complex genome, transcriptome, and predicted proteome landscape underpinning genetic diversity. Mol. Plant 15, 520–536.

Kersey, P. J. (2019). Plant genome sequences: past, present, future. Curr. Opin. Plant Biol. 48, 1–8.

Li, H. (2018). Minimap2: pairwise alignment for nucleotide sequences. Bioinformatics 34, 3094–3100.

Petek, M., Zagorščak, M., Ramšak, ž., Sanders, S., Tomaž, Š., Tseng, E., et al. (2020). Cultivar-specific transcriptome and pan-transcriptome reconstruction of tetraploid potato. Sci Data 7, 249.

Pham, G. M., Hamilton, J. P., Wood, J. C., Burke, J. T., Zhao, H., Vaillancourt, B., et al. (2020). Construction of a chromosome-scale long-read reference genome assembly for potato. Gigascience 9, giaa100.

Potato Genome Sequencing Consortium, Xu, X., Pan, S., Cheng, S., Zhang, B., Mu, D., et al. (2011). Genome sequence and analysis of the tuber crop potato. Nature 475, 189–195.

Quinlan, A. R., and Hall, I. M. (2010). BEDTools: a flexible suite of utilities for comparing genomic features. Bioinformatics 26, 841–842.

Shumate, A., and Salzberg, S. L. (2020). Liftoff: accurate mapping of gene annotations. Bioinformatics 37, 1639–1343.

Tang, D., Jia, Y., Zhang, J., Li, H., Cheng, L., Wang, P., et al. (2022). Genome evolution and diversity of wild and cultivated potatoes. Nature 606, 535–541.

Tomato Genome Consortium (2012). The tomato genome sequence provides insights into fleshy fruit evolution. Nature 485, 635–641.

Tomaž, Š., Petek, M., Lukan, T., Pogačar, K., Stare, K., Teixeira Prates, E., et al. (2023). A mini-TGA protein modulates gene expression through heterogeneous association with transcription factors. Plant Physiol. 191, 1934–1952.

Yandell, M., and Ence, D. (2012). A beginner’s guide to eukaryotic genome annotation. Nat. Rev. Genet. 13, 329–342.

Yang, X., Zhang, L., Guo, X., Xu, J., Zhang, K., Yang, Y., et al. (2023). The gap-free potato genome assembly reveals large tandem gene clusters of agronomical importance in highly repeated genomic regions. Mol. Plant 16, 314–317.

