## Supplementary Material for "Evidence based Unification of poTato gene models with UniTato collaborative genome browser"

**Contents**

Supplementary figures … p2 - p4
Supplementary tables … p5 - p7

### Supplementary figures

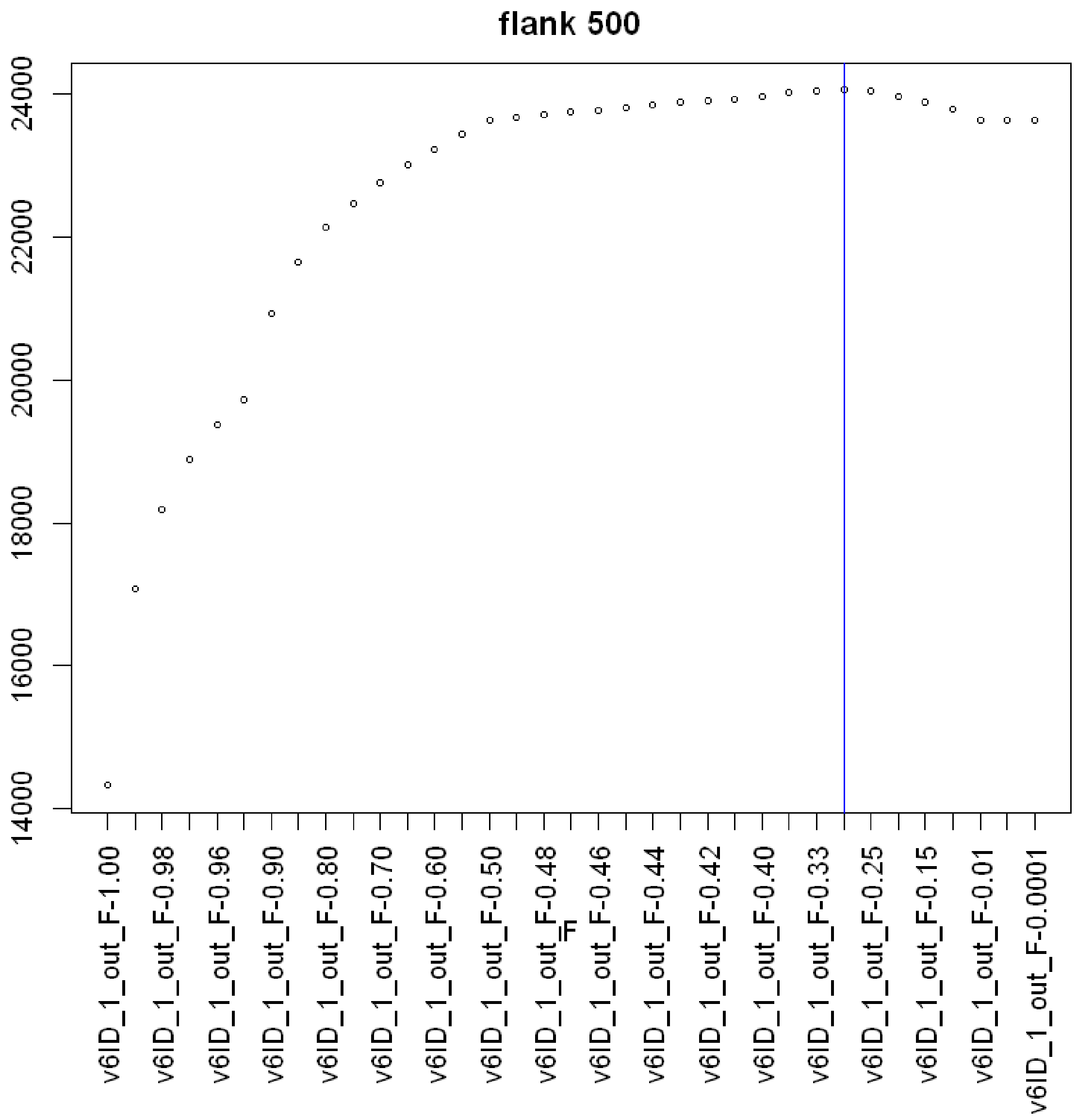


**Figure S1.** Bedtools optimal F threshold selection using ITAG/PGSC pairs mapped to v6 genome with Liftoff (with *flank* = 500) defined in the merged v4n gene model (Petek et al. 2020). The graph shows the number of v4 ITAG/PGSC pairs mapped to the same v6 gene model at decreasing Bedtools F values.

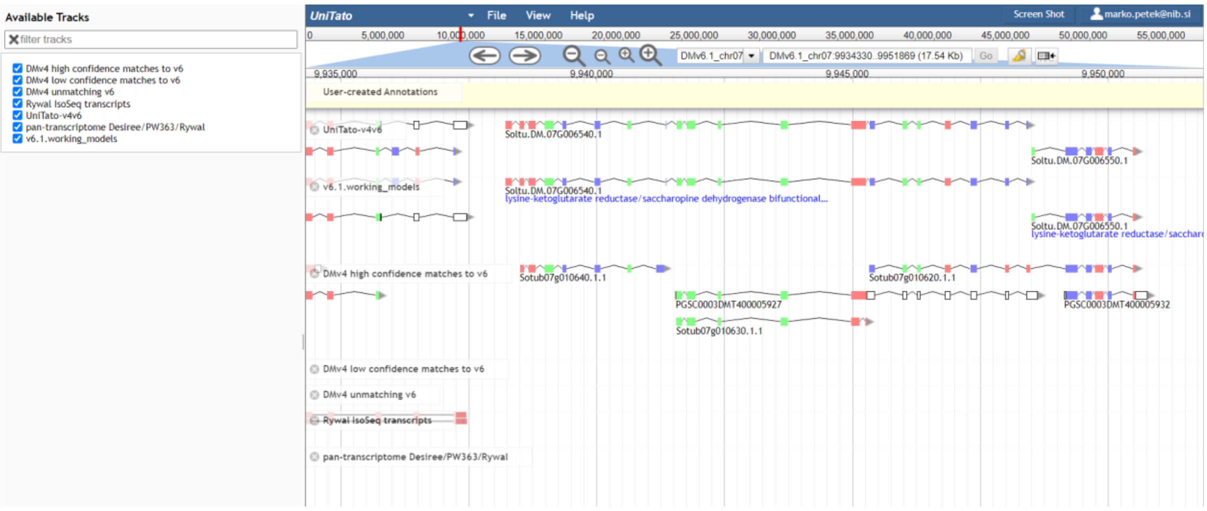


**Figure S2.** An example of v4 and v6 gene models that in our opinion cannot be simply resolved and requires manual curation, possibly based on more evidence. For a list of such gene models see overlaps.xlsx on the Unitato GitHub page (github.com/NIB-SI/unitato).

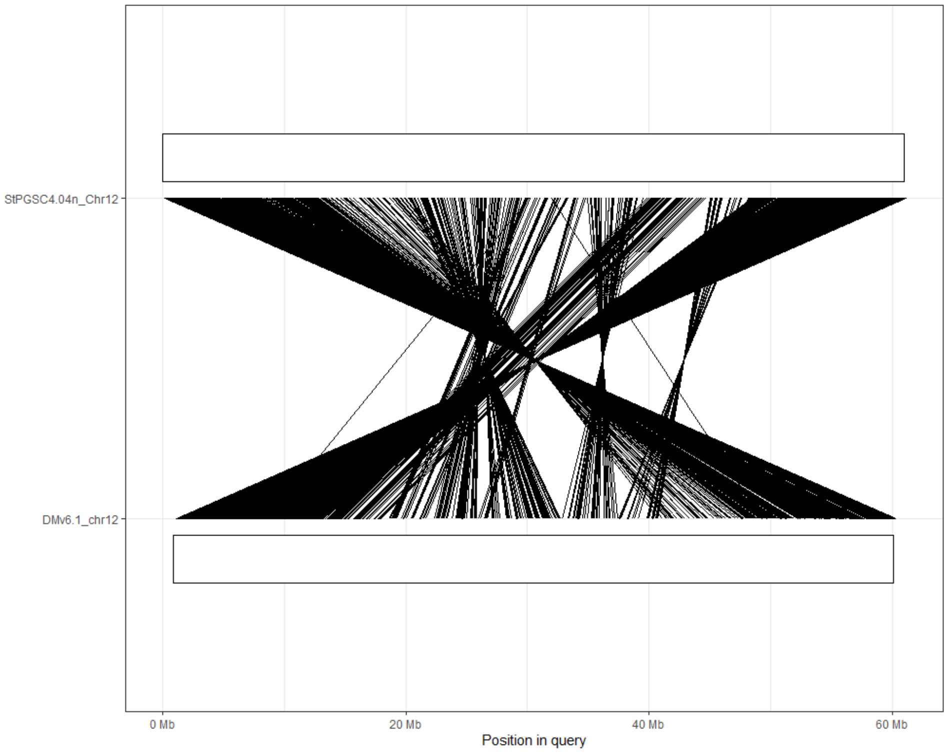


**Figure S3.** Chromosomal 12 rearrangements in v6 genome assembly vs v4 genome assembly. The lines represent synteny between gene model coding regions. Other chromosomes' pairwise synteny graphs can be found on the Unitato GitHub page (github.com/NIB-SI/unitato).

### Supplementary tables

**Table S1.** Overview of total gene counts and Liftoff results at different *flank* parameter values for v4 and v6 gene models. hc - gene models defined as “high confidence” in v6. Note: 316 PGSC and 211 ITAG gene models could not be mapped to the v6 genome assembly (unmapped) with both flank parameter values.

| | Total gene count | No *flank,* unmapped | *Flank* 500 nt, unmapped | No *flank,* mapped | *Flank* 500 nt, mapped |
|---|---|---|---|---|---|
| PGSC | 39,428 (100.00%) | 492 (1.25%) | 363 (0.92%) | 38,936 (98.75%) | 39,065 (99.08%) |
| ITAG | 35,004 (100.00%) | 1,797 (5.13%) | 249 (0.71%) | 33,207 (94.87%) | 34,755 (99.29%) |
| V6.1 working version | 40,652 (32,917 hc) | / | / | / | / |

**Table S2.** Coverage of v4 to v6 gene models by number of models and % of all v6 models, at different Bedtools intersect sequence coverage (*F*) parameter values. Note: genes that mapped with the same F score with and without flank are counted twice.

| | Version | *F* = 1 | *F* >= 0.30 | *F* >= 0.0001 |
|---|---|---|---|---|
| PGSC/ITAG no *flank*, working version | v4 | 40,252 (54.08%) | 56,512 (75.92%) | 57,663 (77.47%) |
| | v6 | 27,103 (66.67%) | 31,481 (77.44%) | 32,263 (79.36%) |
| PGSC/ITAG no *flank*, high-confidence | v4 | 38,095 (51.18%) | 53,360 (71.69%) | 54,199 (72.81%) |
| | v6 | 25,331 (76.95%) | 28,986 (88.06%) | 29,470 (89.53%) |
| PGSC/ITAG *flank* 500 nt, working version | v4 | 40,586 (54.53%) | 57,040 (76.63%) | 58,209 (78.20%) |
| | v6 | 27,261 (67.06%) | 31,669 (77.90%) | 32,452 (79.83%) |
| PGSC/ITAG *flank* 500 nt, high-confidence | v4 | 38,373 (51.55%) | 53,793 (72.27%) | 54,637 (73.40%) |
| | v6 | 25,443 (77.29%) | 29,113 (88.44%) | 29,586 (89.88%) |

**Table S3.** Flank preference in UniTato-v4v6 GFF generation. Mapped genome models with Bedtools coverage F < 0.30 were added to the v6.1 working model GFF3, thus defining UniTato-v4v6. Note: in most cases, genome models from mappings with flank 500 nt were selected. Considering both mappings, without flank and using flank 500 nt, only 527 v4 genome models did not map to the v6, from which 316 were from the PGSC and 211 were from the ITAG annotation.

| | ITAG/PGSCv4 | DMv6 | DMv6 high-confidence |
|---|---|---|---|
| Total gene model count | 74432 | 40652 | 32917 |
| F >= 0.30 flank 0 | 387 | 458 | 368 |
| F >= 0.30 flank 500 | 56776 | 31594 | 29065 |
| 0.0001 <= F < 0.30 flank 0 | 13 | 19 | 14 |
| 0.0001 <= F < 0.30 flank 500 | 1142 | 1418 | 1028 |
| 0 <= F < 0.0001 flank 0 | 32 | / | / |
| 0 <= F < 0.0001 flank 500 | 15555 | / | / |
| unmapped | 527 | / | / |
